# Rapid cost-effective viral genome sequencing by V-seq

**DOI:** 10.1101/2020.08.15.252510

**Authors:** Longhua Guo, James Boocock, Jacob M. Tome, Sukantha Chandrasekaran, Evann E. Hilt, Yi Zhang, Laila Sathe, Xinmin Li, Chongyuan Luo, Sriram Kosuri, Jay A. Shendure, Valerie A. Arboleda, Jonathan Flint, Eleazar Eskin, Omai B. Garner, Shangxin Yang, Joshua S. Bloom, Leonid Kruglyak, Yi Yin

## Abstract

Conventional methods for viral genome sequencing largely use metatranscriptomic approaches or, alternatively, enrich for viral genomes by amplicon sequencing with virus-specific PCR or hybridization-based capture. These existing methods are costly, require extensive sample handling time, and have limited throughput. Here, we describe V-seq, an inexpensive, fast, and scalable method that performs targeted viral genome sequencing by multiplexing virus-specific primers at the cDNA synthesis step. We designed densely tiled reverse transcription (RT) primers across the SARS-CoV-2 genome, with a subset of hexamers at the 3’ end to minimize mis-priming from the abundant human rRNA repeats and human RNA PolII transcriptome. We found that overlapping RT primers do not interfere, but rather act in concert to improve viral genome coverage in samples with low viral load. We provide a path to optimize V-seq with SARS-CoV-2 as an example. We anticipate that V-seq can be applied to investigate genome evolution and track outbreaks of RNA viruses in a cost-effective manner. More broadly, the multiplexed RT approach by V-seq can be generalized to other applications of targeted RNA sequencing.

## Introduction

One approach to obtaining the genome sequences of RNA viruses is to sequence all the RNA molecules present in a sample, including those arising from the virus(es) of interest, other microbes, and the host. Such metatranscriptomic sequencing typically involves depleting abundant human ribosomal RNA (rRNA), which would otherwise account for 90-99% of sequencing reads. An alternative approach is to enrich for the viral genome of interest, which lowers cost by reducing the total amount of sequencing required to obtain a sufficient number of viral reads. Two methodologies for such targeted sequencing are commonly employed: hybridization-based capture with probes that target the viral genome (1) or PCR-based amplicon sequencing. Both of these methods have been used to sequence SARS-CoV-2 genomes (2–6). Metatranscriptomic and targeted approaches share the same first step—converting RNA to cDNA via random-hexamer-primed RT. A Not-So-Random (NSR) hexamer design has been used to eliminate human rRNA during cDNA synthesis; this design, rather than including all 4,096 possible hexamers, uses a subset of 749 that are depleted in rRNA genes (7) and has been employed for analyzing transcriptomes of plasmodium (8). However, NSR-seq does not deplete the human RNA PolII transcriptome, and is therefore not ideal for viral sequencing in samples that also contain human coding transcripts, which greatly exceed viral RNA in total abundance.

To address these limitations, we developed V-seq, a targeted sequencing method that uses virus-specific RT primers tiled across the SARS-CoV-2 genome to enrich for viral sequences (**Fig.1A**). RT-based enrichment is usually less stringent than that by PCR or hybridization because of mis-priming by the 3’ hexamer. We therefore adopted the NSR approach in the design of the 3’ terminal region of the 25 nt RT primers. Further, in order to deplete the human RNA PolII transcriptome, we selected primers with 3’ hexamers that are underrepresented in the human PRO-seq dataset, which encompasses human RNA PolII transcriptome (9). Enrichment by use of single-ended RT primers, although promiscuous, can accommodate highly degraded clinical samples, because the method, unlike PCR, does not require both ends to be present in the same amplicon. Additionally, although we can design overlapping PCR primers in amplicon sequencing, they need to be separated into multiple non-overlapping pools in each PCR reaction, because otherwise shorter overlapping products will be preferably amplified. The number of tiling primers is thus limited. With V-seq, as long as the RT primers do not interfere with each other, they can be densely tiled across the genome in a single pool to improve coverage (**Fig.1B**). Reverse transcriptase enzymes have active RNA-DNA hybrid strand displacement activity (10). With the use of overlapping RT primers, each basepair of the viral transcriptome has multiple opportunities to be converted to cDNA during a single reverse transcription step. We tested this approach and found that overlapping RT primers improve recovery of the viral genome without sacrificing specificity.

The resulting V-seq method has major advantages over current alternatives because the protocol is 1) short (sequencing libraries are generated from extracted RNA in < 5 hours) as it removes the lengthy steps of rRNA depletion, targeted PCR or probe hybridization steps, and simplifies steps for second-strand synthesis (SSS) and addition of indexed sequencing adaptors; 2) scalable to thousands of samples, with an inexpensive cost of $6 per sample (**Table S1**); and 3) generalizable to other applications of targeted RNA-seq. We evaluated different RT primer designs and used V-seq to obtain SARS-CoV-2 sequences from nasopharyngeal (NP) swab samples obtained from COVID-19-positive patients. The performance of V-seq matches conventional methods but, importantly, achieves substantial improvements in library preparation time and costs, which could potentially enable real-time contact tracing during pandemic outbreaks and broader applications in targeted RNA-seq.

**Fig. 1.**
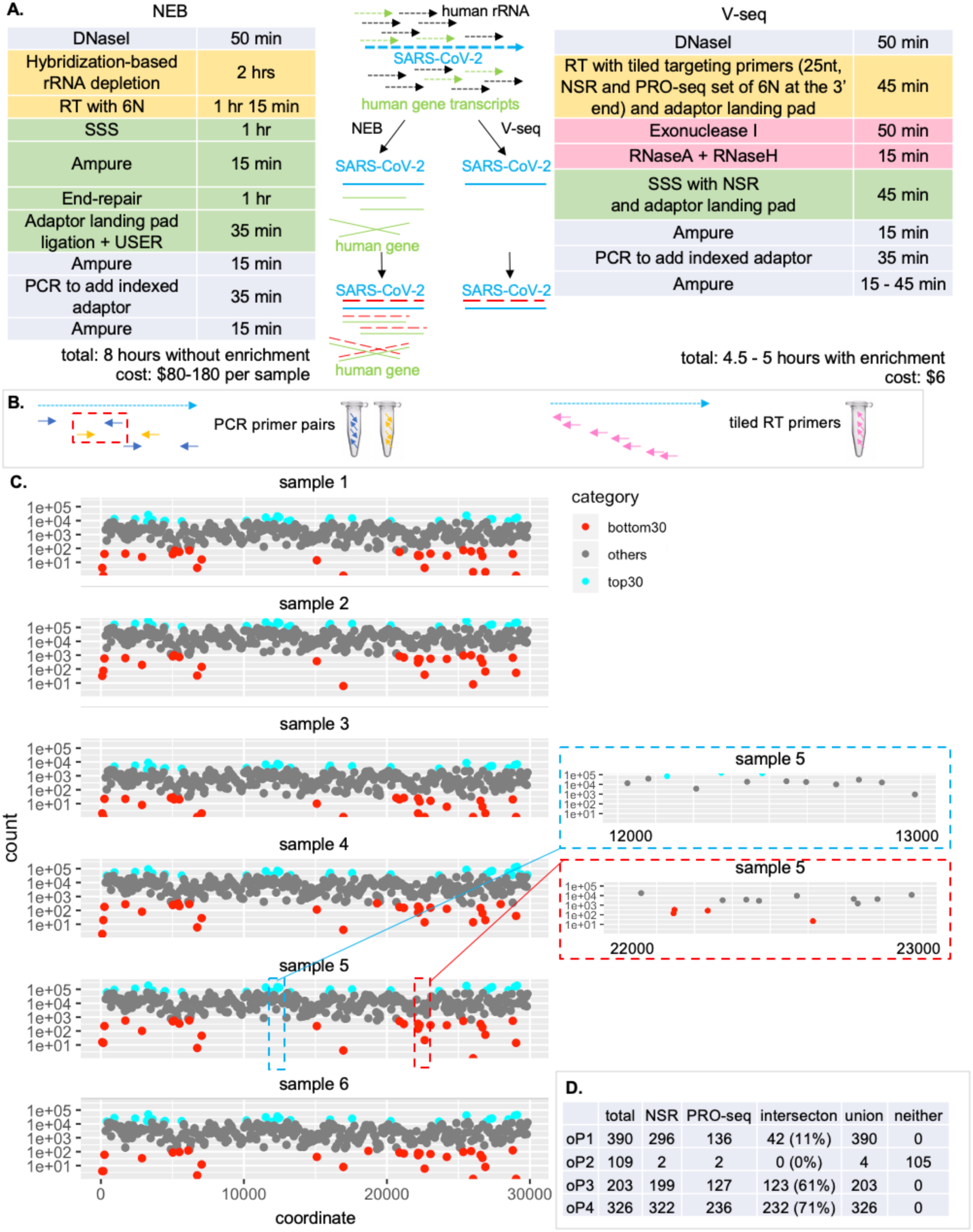
Design of V-seq. **A**. V-seq workflow compared to commercial NEB kit. Dotted arrows are RNA templates. Solid lines are first-strand cDNA. Red dashed lines are second-strand cDNA. Grey shaded boxes are shared steps between NEB and V-seq. Other matching colors show equivalent steps that serve the same purpose but use different methodologies. Adding a PCR-based or hybridization-based enrichment step with the NEB kit takes an additional 2-24 hour processing time. Per sample cost with the NEB kit is roughly $80 without enrichment, and $80-180 with enrichment depending on the methodology and the number of samples multiplexed in the enrichment reaction. **B**. Multiplexed RT primers can be used in one pool while overlapping paired PCR primers need to be separated into different pools limiting the number of total primers one can use to tile across the genome. In amplicon sequencing, e.g., the ARTIC design of SARS-CoV-2 sequencing, overlapping primers are used in two pools (blue and orange) to avoid preferential amplification of short PCR products. Each primer pair generates 300-400 bp amplicons, however, if blue and orange pools are combined, amplification will be preferably between the orange forward primer and the blue reverse primer in the red dashed box. Densely tiled, overlapping RT primers can be used in one pool (pink, right). **C**. Per primer depth across SARS-CoV-2. Left: six samples showing consistent per primer depth. Priming sites with the top 30 and bottom 30 depths are highlighted in blue and red, respectively, which are also consistent across 6 independent samples using oP1 (details in **D**), suggesting reproducibly varied priming efficiency. Right: zoom in on two different regions with relatively high (top) and low (bottom) depth. **D**. Design of the four primer pools. We show the number of total primers in each pool, the number of primers with 3’ hexamers in the NSR or PRO-seq hexamer sets as well as their intersection or union. oP2 corresponds to the ARTIC Network version 3 pool that does not have special design at the 3’ end and thus are largely outside of the NSR/PRO-seq hexamer sets.

## Results

### Overview of V-seq protocol

V-seq uses virus-specific RT primers densely tiled across the viral genome for target enrichment. In **Fig.1A**, we show a comparison between protocols for the commercial NEB kit (see Methods, referred to as “NEB” hereafter) and V-seq. Both methods start with extracted RNA. V-seq removes several time-consuming steps in NEB aimed at rRNA depletion and/or viral enrichment, and achieves the same goal by viral-specific RT with the NSR design. We further eliminate the lengthy end repair, A-tailing and ligation steps in NEB by using an NSR hexamer-primed one-step extension for second strand synthesis, which not only further minimizes rRNA contamination, but also enables adding indexed sequencing adaptors directly by PCR. The resulting protocol can be completed within 5 hours and easily scales to 96-plex indexed libraries, and can potentially be used to multiplex thousands of samples with automation. The key parameter to optimize in V-seq is the design of RT primers.

### Primer design

To overcome challenges of mis-priming (usually initiated from the 3’ end of the RT oligo) from the highly abundant human rRNA and RNA PolII transcriptome, we modified the NSR design (7) as follows: 1) we extracted all unique 25-mers in the reverse complement sequence of the SARS-CoV-2 genome as candidate RT primers; 2) we filtered by GC content, retaining sequences with GC between 40% and 60%; 3) we retained only sequences with a 3’ hexamer that is either in the NSR-seq first strand hexamer set or in the bottom quartile of hexamer abundance in the human PRO-seq dataset (9) (subsequently referred to as NSR and PRO-seq hexamers, **Table S2**); 4) we removed perfect matches to the human genome; and 5) we picked a subset of evenly spaced sequences across the SARS-CoV-2 genome.

We tested four pools of RT primers (**Table S3-6;** see also **Table** S7 for complete oligo sequences for V-seq), as well as specific combinations of these pools. Initially, we designed a 390-primer pool with a median genome spacing between primers of 77 nt (“oP1”) using the criteria above. For comparison, we used the 109 ARTIC Network version 3 reverse PCR primers (2) (“oP2”), which did not use the NSR or the PRO-seq design. After initial experiments with oP1, we found that certain regions of the viral genome were reproducibly sequenced to lower depth (**Fig.1C**). We designed a set of 203 primers, “oP3”, to target these poorly-covered regions, as well as regions with spacing greater than 150 nt between adjacent primers in oP1. We combined oP1 and oP3 to generate the set “oP13”, which has a median spacing of 48 nt between adjacent primer pairs. Lastly, we designed a separate “oP4” set that includes all viral 25-mers such that their 3’ hexamer sequence is present in both the NSR and the PRO-seq hexamer sets, subject to a minimum spacing of 25 nt between primers. We also added 94 primers with 3’ hexamer sequences in either the NSR or the PRO-seq hexamer sets to fill in regions of the viral genome where the gaps between adjacent primers were greater than 182 nt. In total, oP4 contains 326 primers and is expected to most strongly enrich for viral reads, as most of the primers avoid both human rRNA and RNA PolII transcriptome (**Fig.1D**). Each primer pool was synthesized individually at equimolar concentration per primer, and pools were normalized to the same molar concentration per primer when combined.

### Genome coverage and primer specificity as a function of Ct and primer pool combinations

We have sequenced a total of 859M and 1.72B reads for 122 and 138 samples by V-seq and NEB, respectively. These clinical samples were previously tested positive with RT-qPCR assays for SARS-CoV-2, using primers targeting various genes including N, S, and ORF1. In **Fig.2A**, we show viral genome coverage with at least 5x depth as a function of Ct (cycle threshold) from the RT-qPCR test and primer pool combinations. With 90% coverage as the cutoff, we recovered 94 and 71 successful genomes with the median depth of 754x (7M raw reads / sample) and 13x (12.5M) by V-seq and NEB, respectively. The contrast in total raw reads and median depth reflects an average of 90-fold enrichment of viral reads with V-seq compared to NEB, with the caveat being that we have attempted sequencing more high Ct samples with NEB. The average fold enrichment is 87 when we only use samples with Ct less than 25 for both methods. We observed an increase in the fold enrichment of viral reads as Ct increases for samples with Ct less than 25 (**Fig.2B-C**).

**Fig. 2.**
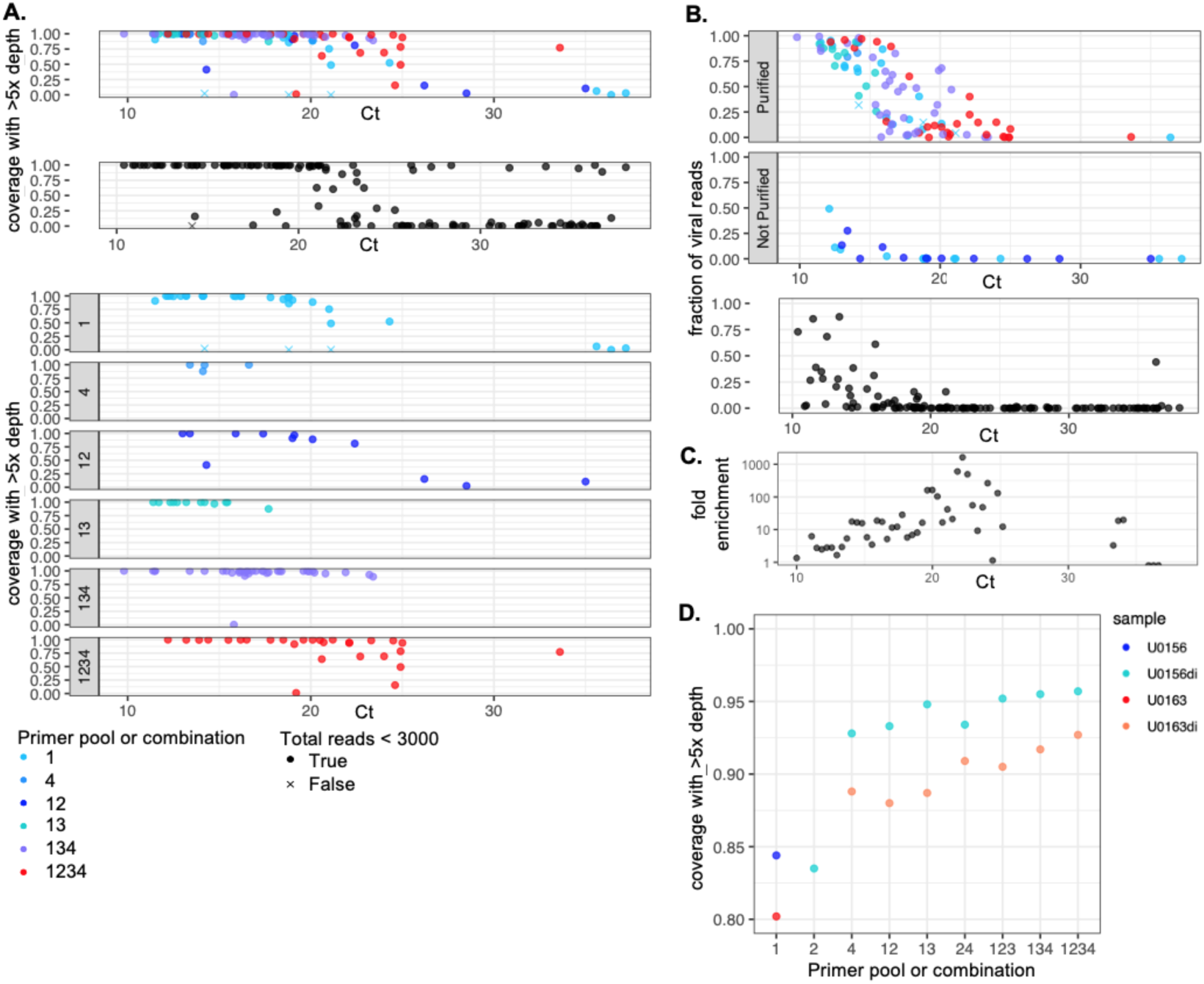
Genome coverage and primer specificity of V-seq as a function of Ct. **A**. Percent of genome covered with at least 5x depth by V-seq (top, colored) and by NEB (top, dark grey) as a function of Ct, as well as breaking down by primer pool (combinations) used (bottom, see **1D** and text for details on primer pools). **B**. Percentage of on-target viral reads as a function of Ct. Colored plots are by V-seq with (top) or without (bottom) a purification step immediately after RT to avoid dimers between RT and SSS oligos (details in Methods). Dark grey plot is by NEB. **C**. Fold enrichment between samples by V-seq over samples by NEB in a sliding window of Ct range of 1. **D**. Genome coverage evaluated on the two sample dilutions down-sampled to 30,000 reads to compare performance of different primer pools. U0156 and U0163 originally have Ct of 12.3 and 13.2, respectively, and were used at 1:10 dilution where “di” was specified.

Both methods robustly work for samples of Ct below 24 with >90% success. Such Ct cutoff covers 2/3 of all the positive samples at UCLA Microbiology Clinical Lab. For the rest of the samples with Ct greater than 24, we had sporadic success (16% of the samples) with NEB in an all-or-none fashion, *i*.*e*., we either obtained >90% coverage or almost entirely failed to obtain any viral reads. With the optimized primer pool combinations, we attempted with V-seq on 5 samples of Ct between 24 and 25 as well as one sample at 33.6, 2 samples had >90% coverage; all the rest had intermediate coverage between 15% and 78%.

In a companion paper, we show that for the 18 shared samples sequenced by both V-seq and NEB, the two methods agree on all the 6,621 variant calls at 380 sites. We trim the primer region of the reads to avoid interfering with variant calling and such regions are excluded in calculating genome coverage, depth, and variant calling [*Guo et al*., personal communication]. In **Fig.S1A**, we compared primers designed with NSR and PRO-seq hexamer ends (oP1, 3, 4 and their combinations) with ARTIC primers (oP2) in an internally controlled manner and found improved performance for our primer pools in terms of genome coverage as a function of Ct. We also observed better on-target specificity. Importantly, denser primers do not seem to interfere with each other but act in concert, which allows us to improve viral genome coverage simply by increasing the number of RT primers.

### Genome coverage and primer specificity evaluated on the same sample dilutions

Results in the clinical samples above suggest that more densely placed primers improve genome coverage, particularly as Ct increases. We further evaluate this trend on the same sample dilutions and down-sampling to the same sequencing depth (**Fig.2D**). We diluted 10-fold for two samples and tested 9 primer pools and/or combinations. Because oP3 was designed to enrich for empirically poorly-covered regions or wide gaps from oP1, we only used oP3 in combination with oP1. Consistent with the trend found in clinical samples, increasing the number of primers improves coverage, particularly for the sample dilutions with higher Ct (“U0163di”). The primer pools that include oP4, which has 3’ hexamers from the intersection of NSR hexamer set and the PRO-seq hexamer set, perform slightly better than the pools without it, as oP4 has the best specificity for viral reads shown in **Fig.S1B** and most effectively excludes contaminating human reads including both rRNA and RNA PolII transcriptome.

### Overlapping RT primers act in concert to improve genome coverage

Improved viral genome coverage by adding additional RT primers suggests that overlapping RT primers do not interfere with each other in their binding to the viral RNA template. In **Fig.3A**, we show positions that are primed more than once by design. We term the primer at the 3’ side in the RT reaction the “preceding” or the “first” primer. In the original oP1 design, 27 out of 390 primers have 3’ end located within 25 nt of the preceding primer. Since the primers are 25 nt in length, we termed these overlapping primer pairs. In oP134, where we added primers in reproducibly poorly-covered regions, 42% of the primers overlap. We then examined all the overlapping RT primers (within 25 nt) and compared their performance with randomly sampled primer pairs from all the non-overlapping primers. As shown in **Fig.1C**, the per primer depth is consistent across samples reflecting their efficiency. Thus, in the analyses below, we use the median depth per primer across all the samples. In **Fig.3B**, we analyzed the ratio of median depth for primers in an overlapping primer pair or in a randomly sampled pair (read counts for the first primer in pair, the preceding primer, divided by read counts for the second primer in the pair). We expect this ratio to be 1 when they do not interfere with each other. As expected, this ratio is not significantly different whether the primer is in overlapping pairs or not (p = 0.73, Mann-Whitney test). In **Fig.3C**, we analyzed the average of median depth between the primers (details in Methods) in an overlapping primer pair or in a randomly sampled pair, and further broke down when the two primers are within 25 or 6 nt. The average efficiency between the two primers is slightly reduced in overlapping pairs (median depth of 635) within 25 nt than in randomly sampled pairs (median depth of 788, p = 0.02, Mann-Whitney test); and not significantly different when evaluated for primers within 6 nt (p=0.66, Mann-Whitney test).

**Fig. 3.**
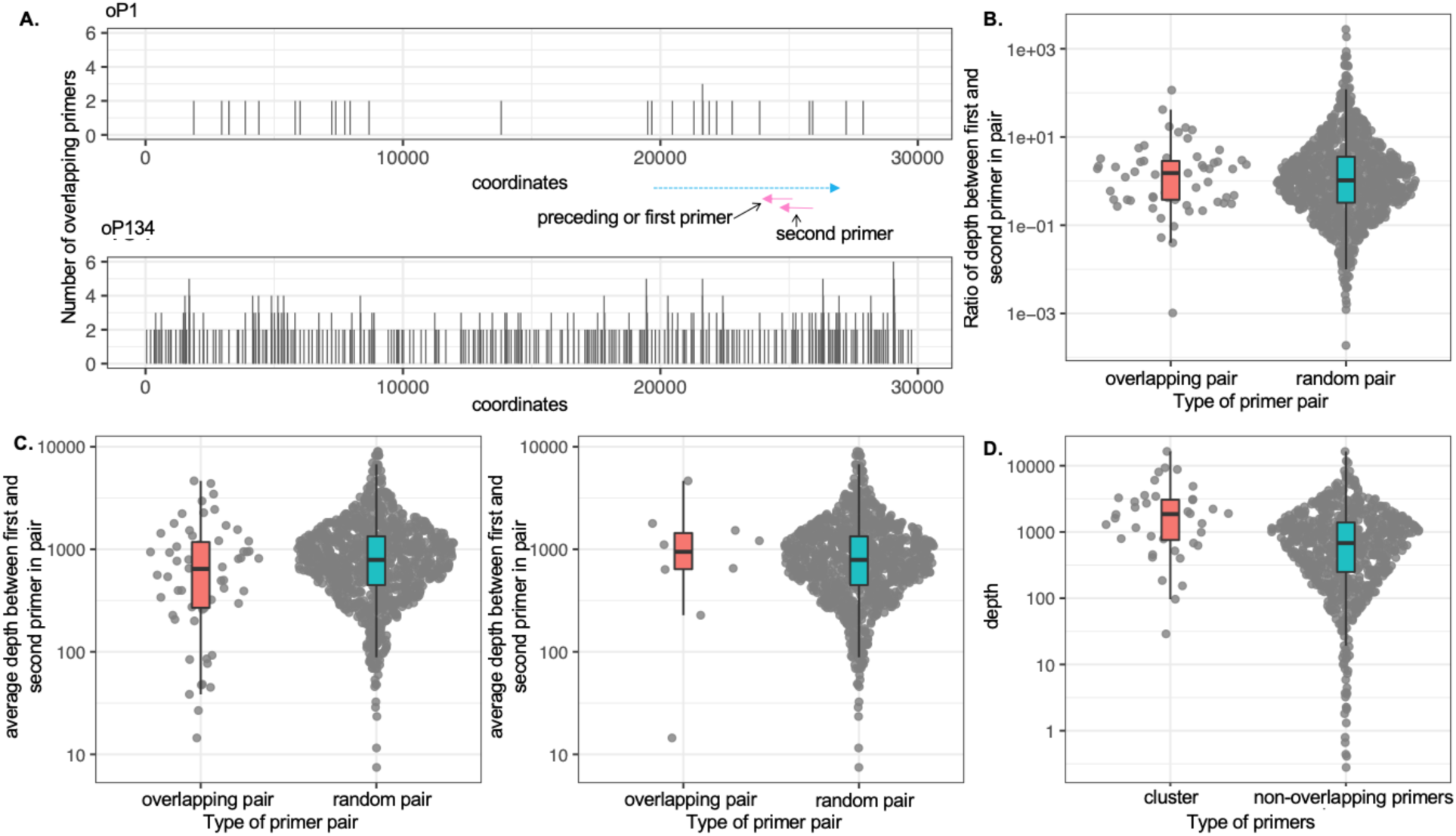
Overlapping primers improve genome coverage in concert. **A**. Repeated coverage by overlapping primers in oP1 (top) and oP134 (bottom). When the 3’ end of two adjacent primers are within 25 nt (pink arrows; blue arrow depicts RNA template), the position of the preceding primer is covered twice and is plotted with count of 2. Occasionally, a third primer runs into the second primer in pair, which is considered as a chain of 3. Note that in oP134, the longest chain involves 6 consecutive primers. **B**. Ratio of depth between the first and the second primers in overlapping pairs or in randomly sampled pairs. If no interference is expected, as in the randomly sampled non-overlapping primer pairs, we would expect this ratio to be centered at 1. The medians of ratios are 1.57 and 1.04 respectively for the overlapping and random pairs (p = 0.73, Mann-Whitney test). **C**. Average depth of the two primers in overlapping pairs or in randomly sampled pairs. Left: overlapping pairs within 25 nt. The median average depths are 635 and 788, respectively, with p-value = 0.02, Mann-Whitney test. Right: overlapping pairs within 6 nt. The median average depths are 957 and 788, respectively, with p-value = 0.66, Mann-Whitney test. **D**. Sum of depths covered by primers in chain (“cluster”) compared to non-overlapping primers, the median depths are 1862 and 674, respectively, with p-value = 6.7e-06, Mann-Whitney test. Note that the min depth in chained primers is improved to 28, while 37 regions were covered with median less than 10 by non-overlapping primers.

Most importantly, we evaluated whether having multiple off-setting primers helps recover otherwise poorly-covered regions. In **Fig.3D**, we show that with overlapping primers, we increased depth for 37 poorly-covered regions (roughly 5% of the genome with depth less than 10 by a single primer) by 2.7-fold (p = 6.7e-06, Mann-Whitney test), with an improved minimal depth of 28. Consistently, we show that the specificity for targeting the viral genome quantified as the percent of on-target reads also does not strongly differ regardless of whether the primer is overlapping or not (**Fig.S2**). However, the preceding primers in the overlapping pair are slightly less specific with decreased percentage of on-target viral reads from 78% to 67% (p = 0.007, Mann-Whitney test). One explanation could be that the strand displacement activity by the RT enzyme from the second primer could promote off-target re-annealing.

## Discussion

NSR-seq was originally developed as a metatranscriptomic approach (7, 8). Recently, a targeted RNA sequencing approach, multiplexed primer extension sequencing (MPE-seq), was developed for studying splicing isoforms (11). MPE-seq uses highly multiplexed RT primers as an alternative strategy to conventional amplicon sequencing or post-PCR hybridization-based methods. The method uses biotin-based pull-down without NSR or PRO-seq designs, and found median enrichment at targeted regions decreases to six fold with increased number of primers. Studying splicing, unlike viral genome sequencing, does not require full coverage of the targeted transcripts. V-seq inherits advantages of both NSR-seq and MPE-seq by selecting primer ends to minimize off-target priming and using highly multiplexed primers to obtain full viral genome coverage.

We show that V-seq is more cost-effective than commercial solutions (a factor of 15 cheaper for library construction) while providing comparable performance (90-fold enrichment of on-target sequencing reads). V-seq robustly calls sequence variants and provides sufficient genome coverage for viral strain analysis even for high Ct samples, for which full genomes cannot be obtained due to low viral load and sample degradation. For sequencing SARS-CoV-2, we recommend using the oP4 primer pool, which has the highest specificity, for low Ct samples (Ct<18), and oP134 or oP1234 for samples with Ct above 18. For designing a primer pool for a new RNA target, we recommend: 1) starting with the method we used for designing oP4, which includes non-overlapping primers with 3’ hexamers that avoid both rRNA and human RNA PolII transcriptome, as well as primers from the union of the two hexamer sets to supplement gaps >200 nt between adjacent primers; 2) performing a V-seq experiment with the candidate pool; and 3) adding overlapping primers in regions with low coverage.

Importantly, the V-seq protocol replaces time-consuming steps from conventional metatranscriptomic or amplicon-based sequencing approaches. This allows for improved throughput and scale, and we estimate that V-seq can multiplex thousands of samples with light automation. In viral outbreaks, viral sequencing can provide capabilities to rapidly trace viral spread and evolution.

V-seq has a number of potential applications beyond SARS-CoV-2 genome sequencing in positive patient samples. Several NGS-based approaches for high-throughput SARS-CoV-2 testing have recently been developed (12). Briefly, these methods use a small number of viral-specific RT primers with sequencing adaptors, followed by detection based on normalized viral read counts. Previous targeted RNA-seq approaches all employ nonspecific random hexamer primed RT, and as a result are incompatible with NGS-based viral detection approaches that require virus-specific RT. V-seq naturally integrates with NGS-based testing and can provide viral strain information concurrently with detection. V-seq can also be leveraged to select highly efficient and specific primers for NGS-based testing, assess their cross-reactivity in a pooled manner, and include primers for other pathogens such as influenza for differential diagnostics.

V-seq can also serve as an optimization platform for other amplicon-based diagnostics and genetic testing, for example, detection of pathogenic non-coding RNAs or splicing errors. Analysis of clinical samples with a wide Ct range provides a useful dataset for optimizing sequencing of targeted transcripts with abundance varying across 4 orders of magnitude. We learned that densely tiled RT primers improve coverage of targeted transcripts, and that overlapping primers do not interfere, but rather act in concert for recovering difficult regions. These results may help to overcome technical difficulties of targeted RNA-seq in applications such as single-cell targeted sequencing of sgRNA in CRISPR screens and of lineage-tracing barcodes. In summary, we anticipate that V-seq is a cost-effect and scalable tool to rapidly investigate genome evolution of RNA viruses and to improve many other applications of targeted RNA sequencing.

## Author Contributions

YY and JMT designed V-seq. LG performed all the experiments. YY, JB, JSB analyzed the data. SC, EEH, SY and OBG provided clinical samples. YZ, LS, XL, CL, VA, JF, EE provided key support of reagent, resource, sequencing and sample acquisition. YY, LK and LG wrote the manuscript with input from JSB, JMT, VA, JB, SY, OBG, SK and all of the authors. YY, LK, JSB supervised the work.

## Acknowledgments

We thank the UCLA David Geffen School of Medicine’s Dean’s Office for their support, the Fast Grants, Inc for funding of this work. A generous donation was provided by Jane Semel. This work was supported by funding from NIH (DP5OD024579 to VA), the Howard Hughes Medical Institute (to LK), and Damon Runyon Cancer Research Foundation (DFS-43-20 to YY).

---

**Fig. S1.**
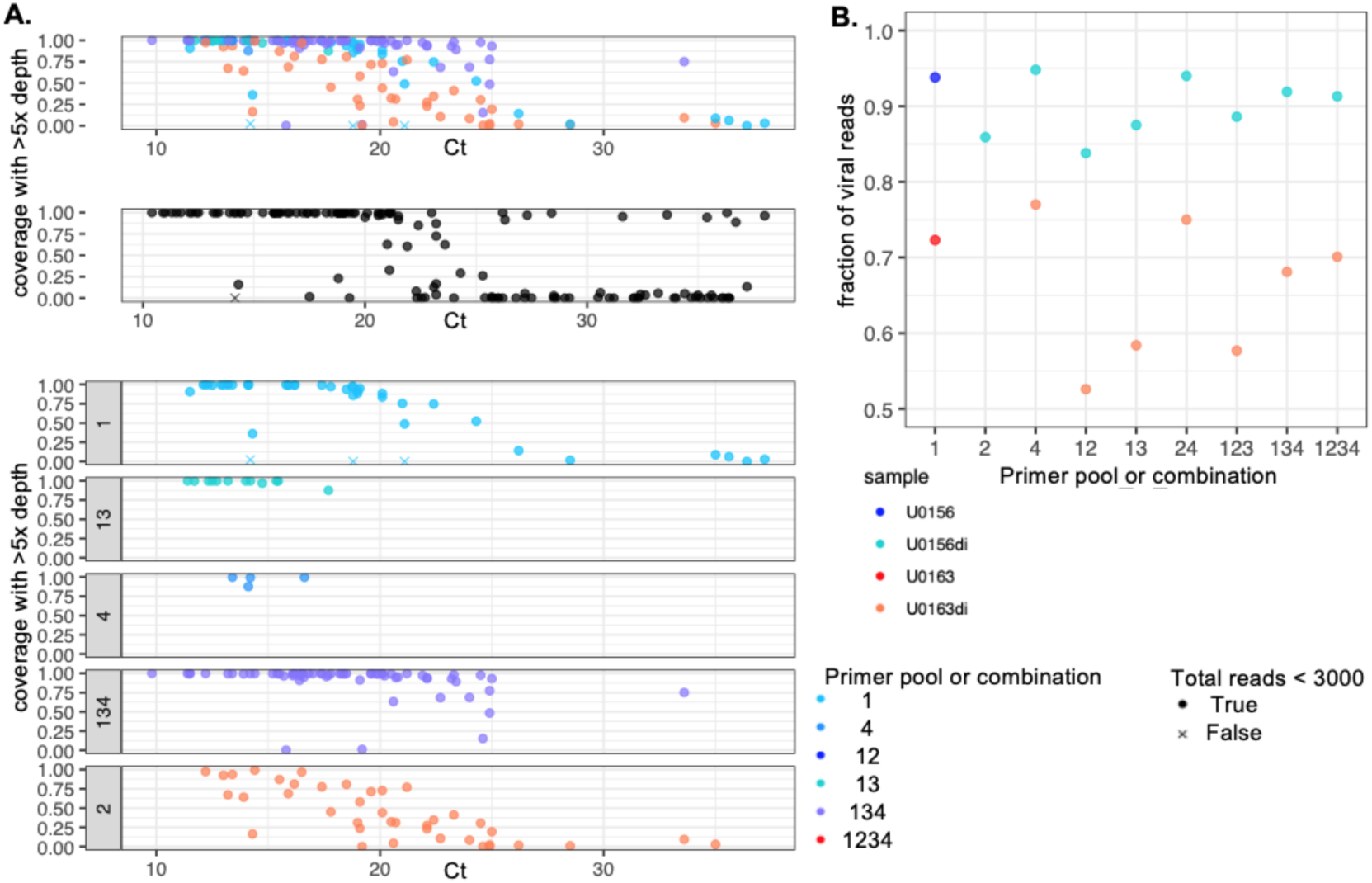
Genome coverage compared to ARTIC design and primer specificity of different V-seq primer pools on the same sample dilutions. **A**. Comparison between V-seq design of primer pool (oP1, 3, 4 and their combinations) with ARTIC V3 design in terms of genome coverage as a function of Ct. **B**. Primer specificity evaluated on the same sample dilution down-sampled to 30,000 reads.

**Fig. S2.**
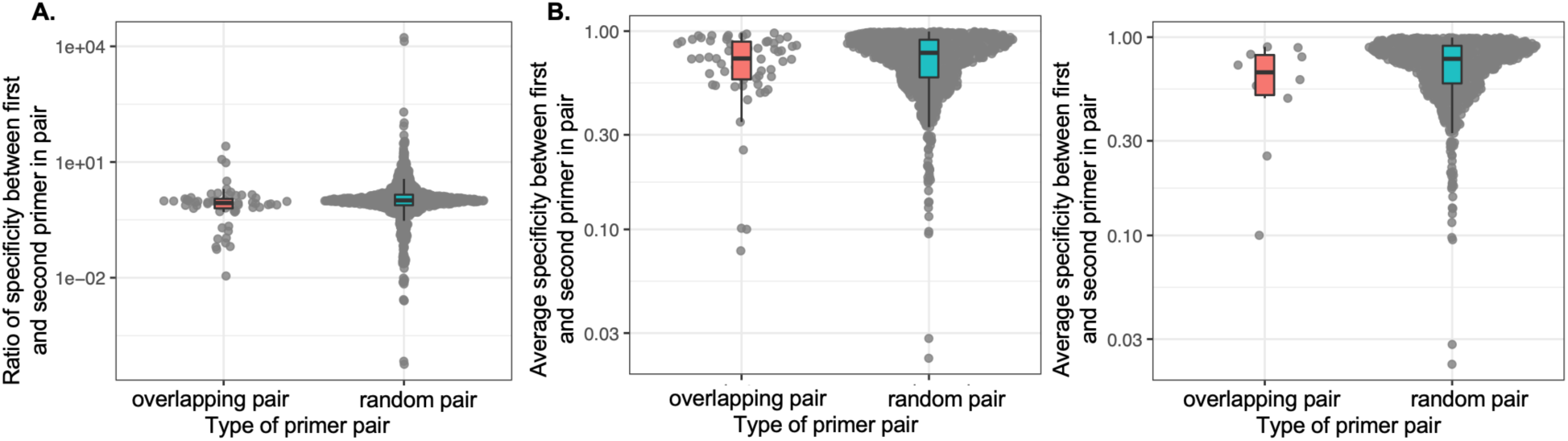
Overlapping primers do not strongly affect primer specificity. **A**. Ratio of specificity (defined as percent of viral reads among raw reads) between the first and the second primers in overlapping pairs or in randomly sampled pairs. If no interference is expected, as in the randomly sampled non-overlapping primer pairs, we would expect this ratio to be centered at 1. The medians of ratios are 0.87 and 1.01 respectively for the overlapping and random pairs (p = 0.007), indicating that the preceding primer is slightly less specific. **B**. Average specificity of the two primers in overlapping pairs or in randomly sampled pairs. Left: overlapping pairs within 25 nt. The median average specificities are 0.67 and 0.78, respectively, with p-value = 0.13, Mann-Whitney test. Right: overlapping pairs within 6 nt. The median average specificities are 0.73 and 0.78, respectively, with p-value = 0.28, Mann-Whitney test.

## References

1. Ng, S.B., Turner, E.H., Robertson, P.D., Flygare, S.D., Bigham, A.W., Lee, C., Shaffer, T., Wong, M., Bhattacharjee, A., Eichler, E.E., et al. (2009) Targeted capture and massively parallel sequencing of 12 human exomes. Nature, 461, 272–276.

2. Itokawa, K., Sekizuka, T., Hashino, M., Tanaka, R. and Kuroda, M. Disentangling primer interactions improves SARS-CoV-2 genome sequencing by the ARTIC Network’s multiplex PCR. bioRxiv, 10.1101/2020.03.10.985150.

3. Wen, S., Sun, C., Zheng, H., Wang, L., Zhang, H., Zou, L., Liu, Z., Du, P., Xu, X., Liang, L., et al. (2020) High-coverage SARS-CoV-2 genome sequences acquired by target capture sequencing. J. Med. Virol., 10.1002/jmv.26116.

4. Xiao, M., Liu, X., Ji, J., Li, M., Li, J., Yang, L., Sun, W., Ren, P., Yang, G., Zhao, J., et al. (2020) Multiple approaches for massively parallel sequencing of SARS-CoV-2 genomes directly from clinical samples. Genome Medicine, 12.

5. Chen, Chen C., Li, J., Di, L., Jing, Q., Du, P., Song, C., Li, J., Li, Q., Cao, Y., et al. MINERVA: A facile strategy for SARS-CoV-2 whole genome deep sequencing of clinical samples. bioRxiv, 10.1101/2020.04.25.060947.

6. Pillay, S., Giandhari, J., Tegally, H., Wilkinson, E., Chimukangara, B., Lessells, R., Moosa, Y., Gazy, I., Fish, M., Singh, L., et al. Whole Genome Sequencing of SARS-CoV-2: Adapting Illumina Protocols for Quick and Accurate Outbreak Investigation During a Pandemic. bioRxiv, 10.1101/2020.06.10.144212.

7. Armour, C.D., Castle, J.C., Chen, R., Babak, T., Loerch, P., Jackson, S., Shah, J.K., Dey, J., Rohl, C.A., Johnson, J.M., et al. (2009) Digital transcriptome profiling using selective hexamer priming for cDNA synthesis. Nat. Methods, 6, 647–649.

8. Vignali, M., Armour, C.D., Chen, J., Morrison, R., Castle, J.C., Biery, M.C., Bouzek, H., Moon, W., Babak, T., Fried, M., et al. (2011) NSR-seq transcriptional profiling enables identification of a gene signature of Plasmodium falciparum parasites infecting children. J. Clin. Invest., 121, 1119–1129.

9. Kwak, H., Fuda, N.J., Core, L.J. and Lis, J.T. (2013) Precise maps of RNA polymerase reveal how promoters direct initiation and pausing. Science, 339, 950–953.

10. Whiting, S.H. and Champoux, J.J. (1998) Properties of strand displacement synthesis by Moloney murine leukemia virus reverse transcriptase: mechanistic implications. J. Mol. Biol., 278, 559–577.

11. Xu, H., Fair, B.J., Dwyer, Z.W., Gildea, M. and Pleiss, J.A. (2019) Detection of splice isoforms and rare intermediates using multiplexed primer extension sequencing. Nat. Methods, 16, 55–58.

12. Bloom, J.S., Jones, E.M., Gasperini, M., Lubock, N.B., Sathe, L., Munugala, C., Sina Booeshaghi, A., Brandenberg, O.F., Guo, L., Simpkins, S.W., et al. Swab-Seq: A high-throughput platform for massively scaled up SARS-CoV-2 testing. medRxiv, 10.1101/2020.08.04.20167874.

