## Supplementary material for "Rapid cost-effective viral genome sequencing by V-seq": Methods

**Supplemental Methods**

**V-seq protocol**

1. Prepare 50uM working stocks for each primer pools (See **Table S7** for oligo sequences; order from IDT with the “oPools Oligo Pools” option).
   oP1: 390 oligos (WGS_oP1_rt)
   oP3: 203 oligos (WGS_oP3_rt)
   oP4: 326 oligos (WGS_oP4_rt)

2. Prepare equimolar primer pool combinations (e.g., 3.9 uLof oP1, 2.03 uL of oP2, 3.26 uL of oP3.

3. Add 1.35uL DNase I enzyme/buffer mix to 7.65uL of purified RNA samples.

| | 1 sample (uL) | Make a master mix of 14 for 12 samples (uL) |
|---|---|---|
| Sample | 7.65 | |
| 10x DNase I buffer | 0.9 | 12.6 |
| DNase I | 0.45 | 6.3 |
| total | 9 | |

   37C 30 min, 75C 20 min

4. Prepare a mix of RT oligos, dNTP and Superase-In.

| | 1 sample (uL) | Make a master mix of 14 for 12 samples (uL) |
|---|---|---|
| primer pool | 0.3 | 4.2 |
| dNTP | 1 | 14 |
| Superase-In | 0.25 | 3.5 |
| total | 1.55 | 21.7 |

5. Add 1.55uL of RT oligo mix from step 4 to the 9uL mix from step 3.
   70C 1 min, 65C 5 min. Hold at 53C.

6. Prepare RT enzyme mix and pre-warm to 53C on a separate thermocycler.

| | 1 sample (uL) | Make 10% more for 12 samples (uL) |
|---|---|---|
| 5x SSIV buffer | 3 | 39.6 |
| 0.1M DTT | 0.75 | 9.9 |
| Superase-In | 0.5 | 6.6 |
| SSIV | 0.5 | 6.6 |
| Total | 4.75 | |

7. Immediately after the sample temperature drops to 53C, add 4.66ul RT enzyme mix per sample while keeping the sample at 53C. Proceed to first strand synthesis.
   53C 15 min, 60C 10 min, 65C 12 min, 70C 8 min, 75C 5 min. Hold at 25C.

8. Add 0.5uL 20U/uL Exonuclease I per sample to remove extra DNA oligos.
   37C 30 min, 80C 20 min.

9. Add 1uL of RNaseA and RNaseH mix per sample.

| | 1 sample (uL) | Make a master mix of 18 for 12 samples (uL) |
|---|---|---|
| RNase A | 0.15 | 2.7 |
| RNase H | 0.25 | 4.5 |

| | | |
|---|---|---|
| Water | 0.6 | 10.8 |
| Total | 1 | |

37C 15 min.

10. Add 1uL of 100uM yy_SSS_R_NTv2_i7 per sample (for better specificity, one can use NSR second strand hexamers at 100uM. The sequences are included in Table S7, see “WGS_SSS”; order from IDT with the “oPools Oligo Pools” option).
    98C 3 min. Leave the samples on bench top to gradually cool down to r.t. (~5min).

11. Add 32.5uL of Kleonow (exo-) mix per sample.

| | 1 sample (uL) | Make a master mix of 13 for 12 samples (uL) |
|---|---|---|
| 10x NEB Buffer 2 | 5 | 65 |
| Klenow (exo-) | 3 | 39 |
| dNTP | 1.8 | 23.4 |
| Water | 22.7 | 295.1 |
| Total | 32.5 | |

25C 15min, 30C 20 min.

12. Add 40uL (0.8x) of Ampure beads per sample. Purify as instructed by the Ampure manual (note that the 80% EtOH needs to be prepared fresh). Elute with 12uL 0.1xTE. Take 10uL to proceed as PCR template.

13. Prepare PCR reactions with dual indexes and template from step 12.

| | 1 sample (uL) | Make a master mix of 13 for 12 (uL) |
|---|---|---|
| Template | 10 | |
| Index 1 | 1.5 | |
| Index 2 | 1.5 | |
| 2x NEBNext mix | 15 | 195 |
| Water | 2 | 26 |
| Total | 30 | |

*Index 1 primers (R2) are included as yy_WGS_P7_1 to yy_WGS_P7_8 in **Table S7**
*Index 2 primers (R1) are included as yy_WGS_P5_1 to yy_WGS_P5_8 in **Table S7**

14. Amplify the genome libraries with the following conditions on a PCR cycler.
    98C 30 sec, (98C 10 sec, 65C 2 min) x 9 cycles, 72C 5 min. 4C hold.

15. Add 30uL of water per sample to bring up the volume to 60uL. Add 48uL (0.8x) of Ampure beads. Purify as instructed by the Ampure manual. Elute with 12uL of 0.1x TE.

16. Optional: depending on the amount of short primer dimers, further Ampure purification may be necessary.

17. NGS requires spiking in Read1 and Index2 primers into existing primer wells. (“yy_WGS_v2_seqR1” and “yy_WGS_v2_seqI2” in **Table S7**)

**Methods with commercial NEB kits**

We followed the manuals for the following NEB kits:

NEBNext® Ultra™ II RNA Library Prep Kit for Illumina® (NEB, E7770L)

NEBNext® Multiplex Oligos for Illumina® (Dual Index Primers Set 1) (NEB, E7600S)

NEBNext® rRNA Depletion Kit (Human/Mouse/Rat) with RNA Sample Purification Beads (NEB, E6350X)
